# WSB1 Regulates c-Myc Expression Through β-catenin Signaling and Forms a Feedforward Circuit Promoting the Development of Cancer

**DOI:** 10.1101/2020.09.25.312678

**Authors:** Xiaomeng Gao, Yanling Gong, Jieqiong You, Meng Yuan, Haiying Zhu, Liang Fang, Hong Zhu, Meidan Ying, Qiaojun He, Bo Yang, Ji Cao

**Author notes:** These authors contributed equally (Xiaomeng Gao, Yanling Gong, Jieqiong You). Corresponding author: Ji Cao, Room 115, College of Pharmaceutical Sciences, Zhejiang University, Hangzhou, China.

## Abstract

The dysregulation of transcription factors is widely associated with tumorigenesis. As the most well-defined transcription factor in multiple types of cancer, c-Myc can directly transform cells by transactivating various downstream genes. Given that there is no effective way to directly inhibit c-Myc, c-Myc targeting strategies based on its regulatory mechanism hold great potential for cancer therapy. In this study, we found that WSB1, a direct target gene of c-Myc, can positively regulate c-Myc expression, which forms a feedforward circuit promoting cancer development. Luciferase-based promoter activity assays and RNA sequencing results confirmed that WSB1 promoted c-Myc expression through the β-catenin pathway. Mechanistically, WSB1 affected β-catenin destruction complex-PPP2CA assembly and E3 ubiquitin ligase adaptor β-TRCP recruitment, which inhibited the ubiquitination of β-catenin and subsequently transactivated c-Myc. Of interest, the promoting effect of WSB1 on c-Myc was independent of its E3 ligase activity. Moreover, co-expression of WSB1 and c-Myc strongly enhanced the initiation and progression of tumours both *in vitro* and *in vivo*. Thus, our findings revealed a novel mechanism involved in tumorigenesis in which the WSB1/c-Myc feedforward circuit played an essential role, highlighting a potential c-Myc intervention strategy in cancer treatment.

## Background

Transcription factors (TFs) recognize DNA in a sequence-specific manner and are responsible for genome transcription control[1, 2]. Given that cancers arise from the abnormal activation of oncogenes or the inactivation of anti-oncogenes, it is not surprising that mutations and dysregulation of TFs are associated with tumourigenesis[3]. Moreover, the dysregulated activity of TFs could in turn cause cancer cells to become highly demanding for certain regulators of gene expression, which represents a common tumour-associated phenomenon called “transcriptional addiction”[4]. To date, hundreds of tumour-associated TFs have been identified. c-Myc is one of the most well-documented tumour-associated TFs in multiple types of cancer and directly transforms cells by transactivating downstream genes involved in cell division, cell cycle progression, angiogenesis, cell senescence and tumour progression[5–7].

Although c-Myc is widely accepted as a promising target for cancer therapy, it lacks a specific ligand binding domain, which limits the development of targeting strategies[8]. Moreover, as c-Myc is located in the nucleus, monoclonal antibody drugs with macromolecules are not able to target it effectively[9, 10]. Thus, there is still no effective way to directly target c-Myc. Recently, many studies have shown that the transactivation of c-Myc is regulated by upstream cytokine signalling, transcription factors and related binding proteins[11], among which the Wnt/β-catenin pathway is the most classical pathway[12]. Upon Wnt signalling activation, the ubiquitination of β-catenin is inhibited, and β-catenin is able to accumulate and translocate into the nucleus to activate the transcription of c-Myc[13, 14]. However, the molecular mechanism of the upstream regulation of c-Myc is not fully understood, which hinders the theoretical development of c-Myc targeting strategies.

WD repeat and SOCS box containing 1 (WSB1) is a member of the Elongin B/C-Cullin2/5-SOCS box protein complex (ESC) ubiquitin ligase, which functions as an adaptor for recognizing substrates for ubiquitination. Increasing evidence suggests that WSB1 is correlated with tumour progression, metastasis and drug resistance[15–17]. Moreover, a recent study discovered that WSB1 promoted the degradation of the DNA damage repair associated protein serine-protein kinase ATM and participated in cell senescence induced by GTPase RAS[18]. In our previous study, we found that WSB1 was the target gene of Hypoxia induced factor 1-alpha (HIF1-α) and could promote the degradation of its substrate Rho GDP dissociation inhibitor 2 (RhoGDI2), leading to metastasis of osteosarcoma[19]. Thus, further exploration of WSB1 and its substrate as well as the regulatory mechanism will not only clarify the important role of WSB1 in tumour processes but also provide new targets for cancer treatment.

Interestingly, based on a literature search and experimental data, we analysed the promoter sequence of WSB1 and found that there was a putative c-Myc binding site, which greatly aroused our interest. In this study, we report that WSB1 is a direct target gene of c-Myc. Moreover, WSB1 could promote the expression of c-Myc through the Wnt/β-catenin pathway, which enhances the cancer-promoting effect induced by c-Myc. Thus, our findings reveal a novel mechanism involved in tumorigenesis in which the WSB1/c-Myc feedforward circuit plays an essential role, highlighting that targeting WSB1 might be a potential method to regulate c-Myc in cancer treatment.

## Methods

### Reagents and antibodies

Proteasome inhibitor MG132, β-catenin inhibitor ICG-001 and XAV939 were purchased from MedChemExpress. Leupeptin, Na_3_VO_4_, pyruvic acid sodium salt and WSB1 antibody (#HPA003293) was purchased from Sigma-Aldrich. Non-essential amino acid and L-glutamine was purchased from Invitrogen (Thermo Fisher Scientific) and Polyethylenimine was purchased from Polysciences. TRIzol, T4 ligase and KOD PCR kit were purchased from Takara Bio. SYBR Green supermixes for real-time PCR was purchased from BIO-RAD. c-Myc (#db1667), HA (#db2603), GAPDH (#db106), Ubiquitin (#db935), GSK3 (#db2953) and CK1 (#db3190) antibodies were purchased from Diagnostic Biosystems. c-Myc (#sc-40) antibody for immunohistochemistry was purchased from Santa Cruz Biotechnology. β-catenin (#9562S), AXIN1 (#2087S) and β-TRCP antibodies (#4394S) were purchased from Cell Signaling Technology. Flag antibody was purchased from GenScript. PPP2CA (I3482-I-AP), Lamin B (66095-I-Ig) antibodies were purchased from Proteintech Group. Wnt 3A conditional medium was generated according to previous report[20].

### Cell culture

Human embryonic kidney (HEK) cell lone 293FT which was purchased from Invitrogen (Thermo Fisher Scientific), human liver hepatocellular carcinoma cell line HepG2, human ovarian cancer cell line A2780 and human colon cancer cell line HCT116 which were purchased from the Shanghai Cell Bank of Chinese Academy of Sciences, were cultured in Dulbecco’s modified Eagle’s medium (Gibco, Thermo Fisher Scientific). Human osteosarcoma cell line KHOS/NP was kindly provided by Dr. Lingtao Wu in University of Southern California, and was cultured in Dulbecco’s modified Eagle’s medium (Gibco, Thermo Fisher Scientific). Human hepatocellular carcinoma cell line Bel-7402 as well as HuH-7, human ovarian cancer cell line OVCAR-8, human osteosarcoma cell line U2OS, human colon cancer cell line SW620 and human lung cancer cell line NCI-H460 which were purchased from the Shanghai Cell Bank of Chinese Academy of Sciences, were cultured in RPMI-1640 (Gibco, Thermo Fisher Scientific). Human non-small cell lung cancer cell line A549 from which was purchased from the Shanghai Cell Bank of Chinese Academy of Sciences, was cultured in Ham ‘s F12 nutrient medium (Gibco, Thermo Fisher Scientific). All cells were cultured at 37°C with 5% CO_2_ atmosphere, and all mediums were supplemented with 10% heat-inactivated fetal bovine serum (Gibco, Thermo Fisher Scientific) as well as 100 IU/mL penicillin and 100 ug/mL streptomycin. All cell lines used were authenticated by STR profiling. The cell lines were monitored for mycoplasma contamination every six months.

### Plasmids construction

The pCDH, pGL4.14, Renilla, pMXs-Hu-N-Myc and pMXs-Hu-L-Myc plasmids were obtained from Addgene. The pCCL-WSB1 and pCCL-c-Myc plasmids were purchased from Genscript, and were re-constructed to pCDH vector with N-terminal FLAG tag. pGL4.14-WSB1 promoter and pGL4.14-c-Myc promoter plasmids were cloning from the genome promoter and re-constructed into pGL4.14 vector. The pGM-TCF/LEF1-luciferase plasmid was purchased from Shanghai Yisheng Biotechnology Co., Ltd. The wild type AXIN1 and its deletion mutants D1-D6 plasmids were constructed into pCDH vector with N-terminal HA tag. CK1, GSK3β, AXIN1, β-TRCP as well as β-catenin plasmids were constructed into pCDNA 3.0 vector with N-terminal HA tag. PPP2CA plasmid was reconstructed into pCDH vector with N-terminal Flag tag. The pCMV-R8.91 and pMD2-VSVG plasmids were kindly provided by Dr. Lingtao Wu in University of Southern California. The lentivirus vector pLKO.1 was purchased from Thermo-Open-Biosystem. The shRNAs of WSB1, β-catenin and AXIN1 were synthesized by Boshang Biotechnology and constructed into pLKO.1 vector, sequences were as follows:

shWSB1-1#:5’-CCGGGAGTTTCTCTCGTATCGTATTCTCGAGAATACGATACGAGAGAAACTCTTTTTG-3’
shWSB1-2#:5’-CCGGGATCGTGAGATTACGTACTATCTCGAGATAGTACGTAATCTCACGATCTTTTTG-3’
shβ-catenin-1#:5’-CCGGCAGATGGTGTCTGCTATTGTACTCGAGTACAATAGCAGACACCATCTGTTTTTG-3’
shβ-catenin-2#:5’-CCGGGCTTGGAATGAGACTGCTGATCTCGAGATCAGCAGTCTCATTCCAAGCTTTTTG-3’
shAXIN1-1#:5’-CCGGCTGGATACCTGCCGACCTTAACTCGAGTTAAGGTCGGCAGGTATCCAGTTTTTG-3’
shAXIN1-2#:5’-CCGGCCGAAAGTACATTCTTGATAACTCGAGTTATCAAGAATGTACTTTCGGTTTTTG-3’.

### Lentivirus transduction

Lentivirus was produced by transfecting 293FT cells with pCMV-R8.91 (packaging vector), pMD2-VSVG (envelope vector) and shRNA plasmids or pCDH plasmids by PEI-40000 with the ratio of 5: 1: 5 in opti-MEM (Gibco, Thermo Fisher Scientific). Virions were collected after 48 h after transfection. Bel-7402, 293FT and H460 cells were transduced with lentivirus companied by polybrene (6 mg/ml) at the MOI of 5-10 to obtain stable cell strains.

### Cell lysate preparation and immunoprecipitation

Cells were collected with pre-cold PBS and lysed with 1% NP-40 buffer (25 mM Tris, 150 mM NaCl, 10% glycerol and 1% Nonidet P-40, pH=8.0) with 0.1 mM PMSF, 0.1 mM Na_3_VO_4_ and 5 ug/mL Leupeptin. After centrifugation at 14000 g, 4 °C for 30 min, the protein quantification was operated by Bicinchoninic acid kit from Yeasen Biotech. Flag affinity beads (Genscript) or HA magnetic beads (Bimake) were mixed with cell lysate at 4°C overnight, and washed for 4 times with washing buffer (25 mM Tris, 300 mM NaCl, 10% glycerol and 0.2% Nonidet P-40, pH=8.0). Cell lysates or the affinity beads were heated at 95°C for 15 min with loading buffer (25 mM Tris-HCl, 2% SDS, 1 mg/mL Bromophenol Blue, 50% glycerol and 5% β-mercaptoethanol, pH=6.8).

### Nuclear and cytoplasmic separation

Cells were collected and gently resuspended with cytoplasmic lysis solution Lysis buffer B (100 mM Tris-HCl, 140 mM NaCl, 1.5 mM MgCl2·6H2O, pH=8.4), and then incubated on ice for 5 minutes before centrifuging at 1000 g at 4 °C for 3 minutes, the supernatant was the cytoplasm part while the precipitate were the nucleus components. Added the prepared detergent (Sodium deoxycholate 3 mg/mL with Tween 40) to Lysis buffer B according to the ratio of 1:10, and then vortexed 2-3 times to wash the nuclei sufficiently. Incubating the tube on ice for 5 minutes before centrifuging at 1000 g at 4°C for 3 minutes for 5 minutes. Then discarded the supernatant, and repeat washing for 5-10 times. Added an appropriate amount of 1% NP40 lysate buffer to the nucleus and put it in liquid nitrogen for 1 min, then took it out and thawed on ice. Repeated the freezing and thawing process twice, and then vortex vigorously before centrifuging at 13200 rpm together with the cytoplasmic components for 30 minutes.

### GST pull down assay

GST and GST-WSB1 expressed in *E. coli* BL21 were purified according to previous report[21] and the purified protein was used freshly. Incubate the whole cell lysate of 293FT and GST or GST-WSB1 protein in 1% NP-40 buffer (25 mM Tris-base, 150 mM NaCl, 10% glycerol and 1% Nonidet P-40, pH=8.0) with protease inhibitor cocktails at 4 °C for 4 h, and subsequently added the glutathione-agarose beads to incubate for another 2 h. Washing the beads for 4 times with washing buffer (25 mM Tris, 300 mM NaCl, 10% glycerol and 0.2% Nonidet P-40, pH=8.0) and resolved the proteins by heating at 95 °C with loading buffer (25 mM Tris-HCl, 2% SDS, 1 mg/mL Bromophenol Blue, 50% glycerol and 5% β-mercaptoethanol, pH=6.8) for western blotting.

### Western blotting

Proteins were separated by 8% or 10% Tris-Glycine-SDS page gels for 1 h and transferred on PVDF membrane (0.45 um, Millipore). Then blocked the membrane with 5% skimmed milk in TBS-T buffer (20 mM Tris-HCl, 150 mM NaCl, 0.1% Tween 20, pH=7.6) for an hour. Diluted the antibodies with TBS-T buffer (Flag, HA and GAPDH are 1:5000, others are 1:1000) and put the membrane into the antibody buffer, room temperature 1 h or 4°C overnight. Washed the membrane with TBS-T for 3 times and incubated the membrane with secondary antibody 1:5000 diluted with 5% skimmed milk in TBS-T for 1 h at room temperature. ECL reagent (NEL103E001EA, PerkinElmer) was applied to catch the chemiluminescence signal using Amersham Imager 600 (GE Healthcare).

### Chromatin immunoprecipitation assay

Chromatin immunoprecipitation assay was performed using a CHIP assay kit from Millpore. Birefly, 293FT cells transfected with pCDH-c-Myc plasmid were collected and subsequently cross-linked with formaldehyde. Chromatin was transferred on the ice with sonic treatment to obtain about 300bp fragments. Anti-IgG or anti-c-Myc antibody were incubated with the chromatin fragments at 4°C overnight, and the immunoprecipitated fragments were detected by PCR amplification using P1, P2 and P3 primers of WSB1 promoter region.

P1 forward: TCAAGACCAGCCTGGCCAATATCGTG
P1 reverse: GCTCTGTCTCCCAGAGCAGAGTG
P2 forward: GATGAAGTAAACATGCCCTACAGTG
P2 reverse: TTCACCCTGTTGGCCAGGCTAGT
P3 forward: GCAATGTTTAGGGTCCACACGAG
P3 reverse: ATCCCACTGTCCTGGCCAAGATG

### RNA extraction, Reverse transcriptional PCR and Quantitative Real-time PCR

RNA was extracted with TRIzol reagent (#9109, Takara) from cultured cells according to according to the manufacturer’s instructions. cDNA was acquired by Transcript one-step gDNA Removal and cDNA Synthesis supermix (#AT311-03, Tansgene Biotech). The quantitative real-time RT-PCR analysis was performed by iTaqTM Universal SYBR Green Supremix (#172-5124, BIO-RAD). The reaction mixtures containing SYBR Green were composed following the manufacturer’s protocol and then CT values were obtained using a qPCR platform (QuantStudio 6 Flex Real-Time PCR System, ThermoFisher Scientific). Real-time PCR data was monitored using QuantStudio 6 Design and Analysis Software Version 2.3. Relatively expression levels were normalized to the internal control β-Actin. Primers used in Quantitative Real time-PCR were as follows:

WSB1 forward: ACTGTGGAGATATAGTCTGGAGTCT
WSB1 reverse: GTAGCAAGAAGTAGCTGATCTTGTC
CTNNB1 forward: GTTCAGTTGCTTGTTCGTGC
CTNNB1 reverse: GTTGTGAACATCCCGAGCTAG
c-Myc forward: GGACCCGCTTCTCTGAAAG
c-Myc reverse: GTCGAGGTCATAGTTCCTGTTG
N-Myc forward: GCGACCACAAGGCCCTCAGTACCTC
N-Myc reverse: AATGTGGTGACAGCCTTGGTGTTGG
L-Myc forward: CTGGAGAGAGCTGTGAGCGACCGG
L-Myc reverse: GAGCAGGCCTGGGTCTTGGGTTCG
β-Actin forward: TCACCCACACTGTGCCCATCTACGA
β-Actin reverse: CAGCGGAACCGCTCATTGCCAATGG

### Dual-luciferase reporter assay

Cells were seeded in 96-well plate with density of 80% and transfected with pCDH-c-Myc and pGL4.14-WSB1 promoter-luciferase plasmids using jetPRIME (Polyplus Transfection), the ratio of which was 1:2. After transfection on H460 cells for 36 h, discarded the supernatant medium and lysed the cells using 1x PLB buffer in the Luciferase reporter kit (Promega) for 15 min. Transferred 20 ul of the cell lysis to the white plate and added 50 ul LAR II per well, then detected the Firefly Fluc value in the Microplate reader. Sulforhodamine B stain was used as an internal control.

pCDH-WSB1 and pGL4.14-c-Myc promoter-luciferase plasmids as well as pCDH-WSB1 and pGL4.14-TCF/LEF1-luciferase plasmids were transfected on H460 and tested with the Luciferase reporter kit (Promega) in the same way. Stop & Glo reagent was used to stop the firefly reaction and start Renilla luciferase reaction at the same time, the valve of which was used as an internal control.

### Cell proliferation assay

Bel-7402 stably overexpressing WSB1 or WSB1 with c-Myc and H460 cells knocking down WSB1 were seeded in 6-well plate with density of 10% respectively. After 2 weeks cells were stained by the sulforhodamine B and photographed by E-Gel Imager (Bio-Rad). The cloning numbers in each group were counted to calculate the inhibition rate.

### Sphere formation assay

The medium of sphere assay was DMEM/F-12 accompanied with penicillin/streptomycin antibiotic, basic fibroblast growth factor (bFGF, 10 ng/mL), human recombinant epidermal growth factor (EGF, 10 ng/mL) and N-2 supplements (1 X). Bel-7402 cells and HuH-7 cells were collected and washed with DMEM/F12 medium to remove the serum, and counted the cells to reseed with the density of 3000 cells per 2 ml medium per well of the ultra-low attachment 6-well plate. Incubated the plate at 37°C with 5% CO_2_ for about 5-8 days when the cells could form and reach the size of 100 μm. Photographs were taken by upright microscope (Olympus).

### Immunofluorescence assay

Cells were seeded in confocal plates and were fixed with 4% paraformaldehyde for about 15 min. Blocked the fixed cells with 1% BSA for 30 min in room temperature and incubated with β-catenin antibody (dilution at 1: 100) at 4°C overnight. Anti-rabbit secondary antibody was performed in room temperature for 1 h and the nuclear was stained with 4’, 6-diamidino-2-phenylindole (DAPI, Sigma) for 1-4 min. Photographs were taken by the confocal laser scanning microscope (Leica).

### Nude mice xenograft tumor model

Five to six-week-old female balb/c nude mice (National Rodent Laboratory Animal Resource, Shanghai, China) were used for experiments. The mice were randomly divided into 8 groups. H460 cells after lentivirus infection were subcutaneous injected into both sides of the mice with the density of 1 × 10^6^ and the weights, tumor numbers as well as the tumor size were recorded until the mice were sacrificed. Tumor size (mm^3^) was measured with Vernier calipers and calculated with formula (W^2^ × L)/2, in which W was the width and L was the length. The animal experiment was approved by the Institutional Animal Care and Use Committee in Zhejiang University, and was carried out following the institutional guidelines.

### Immunohistochemistry

HCC tissue array (BC03119b) was purchased and processed from Alena Biotechnology (Xi’an, China) and photographs were taken by upright microscope (Olympus). Sacrificed the nude mice at the end of the xenograft experiment and dissected the tumor fixed the tissues with 4% paraformaldehyde. Paraffin sections were prepared and the antigen retrieval was applied with EDTA solution (50 mM Tris, 10 mM EDTA, pH=9.0) after dewaxing treatment. β-catenin antibody (1: 100), WSB1 antibody (1: 200) and c-Myc antibody (1: 200) were incubated with the tissues overnight at 4°C. DAB chromogenic kit was used to reaction with the second antibody. Photographs were taken by upright microscope (Olympus).

### RNA-seq

Bel-7402 cells stably overexpressing pCDH, pCDH-WSB1 and pCDH-ΔSOCS by lentivirus transfection were harvested and washed for 3 times with pre-cold PBS. Total RNA of the cells was extracted by TRIzol reagent and sent to Bohao Biotechnology (Shanghai, China) for RNA expression profiling.

### Statistical analysis

Data was presented as the form of means ± SD and unpaired two-tailed Student’s t-test was applied for statistical analysis. The group differences were considered significantly with P values less than 0.05 (*: p < 0.05, **: p < 0.01, and ***: p < 0.001).

## Results

### 1. WSB1 is a target gene of c-Myc and can in turn regulate c-Myc expression

In previous reports, we and other groups proved that WSB1 is the target gene of HIF1-α by HIF1-α directly binding to its −339 bp region[19]. Given that the c-Myc binding E boxes (CACGTG) match the hypoxia response element consensus sequence (A/GCGT), which results in a similar pattern of gene regulation between HIFs and the proto-oncogene c-Myc, and that the WSB1 promoter also contains putative c-Myc binding E boxes in its −340 bp region, we reasonably speculated that WSB1 might be a target gene of c-Myc. To confirm this prediction, we first examined the protein levels of WSB1 and c-Myc in 10 cancer cell lines and found that there was a highly positive correlation between the expression of WSB1 and c-Myc in these cell lines from various tissues (Figure 1A and S1A). Moreover, we applied immunohistochemistry analysis to a tissue microarray with 119 hepatocellular carcinoma (HCC) clinical samples, and the results showed that there was a statistically positive correlation between WSB1 and c-Myc protein expression levels in HCC tissues (Figure 1B and 1C).

**Figure 1.**
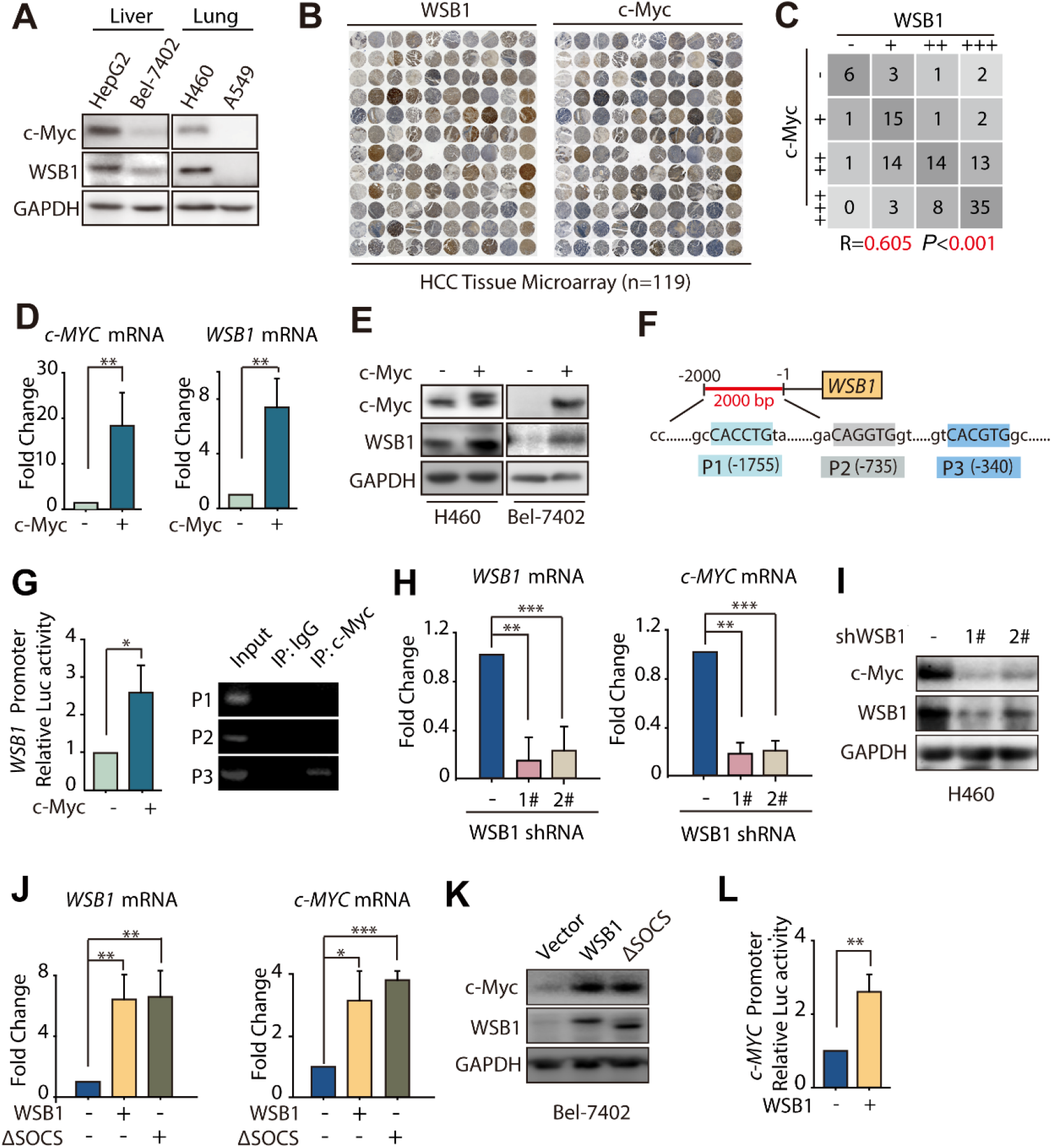
WSB1 is a target gene of c-Myc and could in turn regulate c-Myc expression. (A) Western blotting of WSB1 and c-Myc expression in different tumor cell lines. (B) Immunohistochemical (IHC) staining of WSB1 and c-Myc in hepatocellular carcinoma tissues. (C) Statistical analysis of IHC results of WSB1 and c-Myc in hepatocellular carcinoma tissues (n = 119). (D) Relative mRNA levels of *c-MYC* and *WSB1* in Bel-7402 cells transfected with pCDH-c-Myc plasmid for 48 hours. Data was represented as the means ± SD, n = 3; Statistical significance was determined by Student’s t-test. **, p < 0.01 (versus vector group) (E) Western blotting of WSB1 and c-Myc in H460 and Bel-7402 cells transfected with pCDH-c-Myc plasmid for 48 hours. (F) Schematic representation of the human WSB1 promoter sequence spanning 2.0kb upstream of the transcriptional start site. P1 (−1755), P2(−735) and p3(−340) are predicted potential c-Myc binding sites. (G) Bel-7402 cells were co-transfected with vector or pCDH-c-MYC and pGL4.14-WSB1 promotor for 48 hours. Luciferase activity was detected by M5 and normalized by cell numbers. Data was represented as the means ± SD, n = 3; Statistical significance was determined by Student’s t-test. *, p < 0.05 (versus vector group). 293FT cells were transfected with pCDH-c-Myc plasmid for 48 hours, anti-IgG and anti-c-Myc antibodies were used in the chromatin immunoprecipitation (ChIP) assay. (H) Relative mRNA levels of *c-MYC* and *WSB1* in H460 cells infected with 2 specific lentivirus WSB1 shRNAs for 72 hours. Data was represented as the means ± SD, n = 3; Statistical significance was determined by Student’s t-test. **, p < 0.01 (versus control group). (I) Western blotting of WSB1 and c-Myc in H460 cells infected with lentivirus WSB1 shRNA for 72 hours. (J) Relative mRNA level of *WSB1* and *c-MYC* in Bel-7402 cells infected with lentivirus pCDH or pCDH-WSB1 or pCDH-ΔSOCS for 72 hours. Data was represented as the means ± SD, n = 3; Statistical significance was determined by Student’s t-test. *, p < 0.05; **, p < 0.01; ***, p < 0.001 (versus vector group). (K) Western blotting of WSB1 and c-Myc in Bel-7402 and H460 cells infected with pCDH or pCDH-WSB1 or pCDH-ΔSOCS lentivirus for 72 hours. (L) Bel-7402 cells were co-transfected with vector or pCDH-WSB1 and pGL4.14-c-Myc promotor luciferase for 48 hours. Luciferase activity was normalized by cell numbers and shown as fold change. Data was represented as the means ± SD, n = 3; Statistical significance was determined by Student’s t-test. **, p < 0.01 (versus vector group).

Next, we investigated whether WSB1 was transcriptionally regulated by c-Myc. We overexpressed c-Myc in Bel-7402 cells and subsequently examined both the mRNA and protein levels of *WSB1*. As shown in Figure 1D and 1E, both the mRNA and protein levels were significantly increased upon c-Myc overexpression in Bel-7402 cells. An increase in WSB1 protein expression was also observed in H460 cells with high WSB1 expression under basal conditions (Figure 1E). Additionally, a WSB1 promoter luciferase reporter assay and chromatin immunoprecipitation (ChIP) were performed to confirm whether *WSB1* was the direct target gene of c-Myc (Figure 1F). As shown in Figure 1G, c-Myc overexpression significantly increased WSB1 promoter-based luciferase activity, and the ChIP-PCR results further suggested that c-Myc was able to transcriptionally activate WSB1 expression by directly binding to the −340 bp region of the WSB1 promoter.

We also tested the effect of two other Myc family proteins, N-Myc and L-Myc, on WSB1 expression. The results showed that N-Myc had no effect on WSB1 expression, while L-Myc could promote the mRNA and protein levels of WSB1 (Figure S1B-E). As there is a lack of evidence to show the role of L-Myc in tumourigenesis[22], we thus mainly focused on c-Myc in our study. Collectively, our data indicated that *WSB1* is a target gene of c-Myc.

Encouraged by the fact that c-Myc can transactivate several downstream genes, such as *EBP2*, eIF4F, and *PLK1*, to form feedback loops in tumour progression[23–25], we tested whether WSB1 was able to conversely regulate c-Myc. Notably, decreased c-Myc mRNA and protein levels were detected in H460 cells after WSB1 was knocked down by two specific shRNAs (Figure 1H and 1I). Consistent with these findings, overexpressing wild-type WSB1 in Bel-7402 cells increased both the mRNA and protein levels of c-Myc (Figure 1J and 1K). Moreover, a c-Myc promoter luciferase reporter assay also showed that WSB1 could transactivate c-Myc expression (Figure 1L). Interestingly, when we overexpressed the C-terminal deletion mutant of WSB1 (named ΔSOCS), whose SOCS-box domain was missing and lacked E3 ligase activity, both the mRNA and protein levels of c-Myc were also increased, and comparable effects were observed compared to those of wild-type WSB1 in Bel-7402 cells (Figure 1J and 1K), suggesting that the regulatory effect of WSB1 on c-Myc was independent of its E3 ligase activity. Taken together, all the data mentioned above suggested that WSB1 was a target gene of c-Myc and could in turn regulate c-Myc expression, thus forming a feedforward circuit.

### 2. WSB1 enhances c-Myc expression through the Wnt/β-catenin pathway

Since WSB1 is not a transcription factor, we aimed to decipher the mechanism of how WSB1 promoted the transcriptional level of c-Myc. Thus, RNA-seq was performed to characterize the potential factor in this regulatory process. We stably overexpressed wild-type WSB1 and ΔSOCS WSB1 in Bel-7402 cells and also introduced the vector control by lentivirus infection, and the cells were collected to profile gene expression. RNA-seq results showed that 249 genes were upregulated and 607 genes were downregulated with 2-fold changes out of 18,305 genes in the WSB1-overexpressing group compared to the vector control. Moreover, 252 genes were upregulated and 603 genes were downregulated with 2-fold changes out of 17,936 genes in the ΔSOCS-overexpressing group compared to the vector control (Figure 2A and Table S1). Among the genes with altered expression, 519 genes (139 upregulated genes and 380 downregulated genes) were identified from the two data sets (Figure 2A).

**Figure 2.**
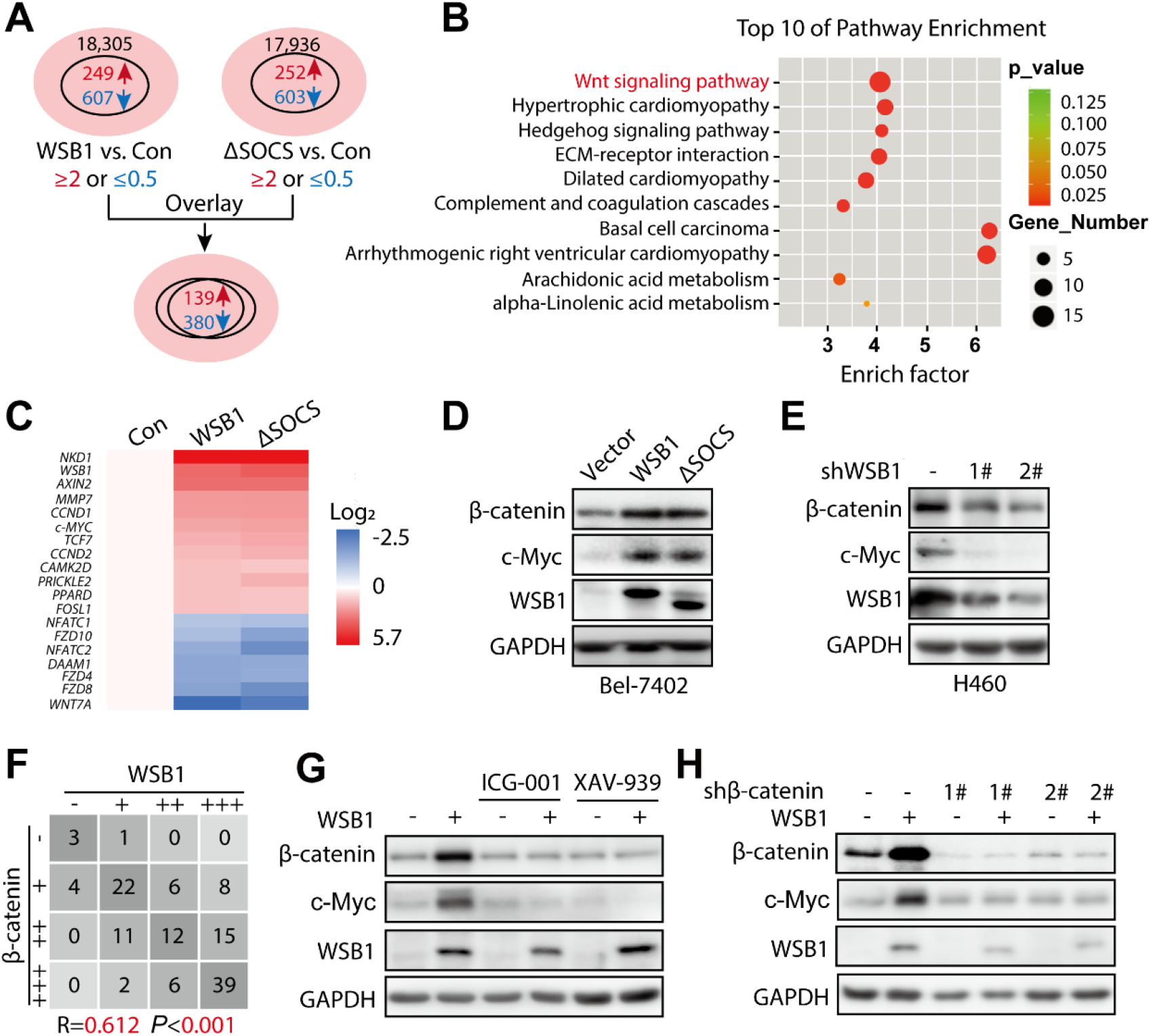
WSB1 enhances c-Myc expression through Wnt/β-catenin pathway. (A) Bel-7402 cells were transduced with pCDH vector or pCDH-WSB1 or pCDH-ΔSOCS for 72 hours. The total RNA were isolated and subsequently performed the RNA-seq. Differential genes (with at least 2-fold change) of WSB1 vs. Control and ΔSOCS vs. Control were overlaid. (B) Differential genes of WSB1 vs. Control and ΔSOCS vs. Control were analyzed by KEGG pathway enrichment analysis and the top 10 pathways were displayed as scatter diagram. (C) Differential genes in Wnt/β-catenin signaling pathway were shown as heatmap. (D) Western blotting of β-catenin, WSB1 and c-Myc in Bel-7402 cells infected with lentivirus pCDH-WSB1 for 72 hours. (E) Western blotting of β-catenin, WSB1 and c-Myc in H460 cells infected with lentivirus WSB1 shRNAs for 72 hours. (F) Statistical analysis of IHC results of WSB1 and β-catenin in hepatocellular carcinoma tissue (n = 119). (G) H460 cells infected with lentivirus Vector or lentivirus pCDH WSB1 were treated with β-catenin inhibitors ICG-001 (10 μM) and XAV-939 (10 μM) for 24 hours. The protein expression of β-catenin, WSB1 and c-Myc were evaluated by Western blotting. (H) H460 cells overexpressing WSB1 or control cells were infected with lentivirus vector or lentivirus β-catenin shRNA for 72 hours. The protein expression of β-catenin, WSB1 and c-Myc were evaluated by Western blotting.

Given that the effect of WSB1 on c-Myc expression was independent of its E3 ligase activity (Figure 1J and 1K), we reasoned that both the wild-type and ΔSOCS WSB1 would activate the factor that regulates c-Myc expression. Thus, KEGG pathway analysis of these 519 overlapping genes was performed, and the results indicated that among the top 10 pathways, arrhythmogenic right ventricular cardiomyopathy, basal cell carcinoma and the Wnt signalling pathway were the top 3 candidates (Figure 2B). Considering that the Wnt/β-catenin signalling pathway is a classical upstream pathway of c-Myc transcription, we performed further analysis of Wnt-regulated gene enrichment. Consistent with the above results, 16 Wnt/β-catenin downstream genes were enriched, including *c-Myc*, that were obviously upregulated or downregulated by WSB1 in our RNA-seq results, indicating the regulation of WSB1 by the Wnt/β-catenin signalling pathway (Figure 2C).

To confirm this finding, we overexpressed WSB1 and ΔSOCS in Bel-7402 cells and found that WSB1 increased the protein levels of both β-catenin and c-Myc (Figure 2D). When we knocked down WSB1 in H460 cells with 2 specific shRNAs, β-catenin and c-Myc levels were downregulated as expected (Figure 2E). Moreover, immunohistochemistry analysis of HCC clinical samples also suggested a highly positive correlation between WSB1 and β-catenin expression (Figure 2F and S2). To prove that WSB1 regulated c-Myc expression through β-catenin, we treated WSB1-overexpressing cells with two pharmacological β-catenin inhibitors, ICG-001 and XAV-939, and subsequently measured β-catenin and c-Myc expression. As shown in Figure 2G, the β-catenin inhibitors blocked the upregulation of both β-catenin and c-Myc expression induced by WSB1. Similar results were also observed when β-catenin was knocked down by lentivirus infection (Figure 2H). Taken together, our data indicated that β-catenin was the key factor participating in WSB1-mediated promotion of c-Myc expression.

### 3. WSB1 promotes β-catenin nuclear translocation and inhibits its ubiquitination

In our previous data, we interestingly found that WSB1 increased the expression of β-catenin (Figure 2D). In line with this, manipulating the WSB1 expression level could also affect the transcriptional activity of β-catenin in the TCF/LEF luciferase reporter assay (Figure 3A and 3B). Under normal conditions without Wnt induction, β-catenin is captured by the destruction complex, subsequently losing its ability to enter the nucleus and is recognized by its E3 ligase adaptor β-TRCP followed by ubiquitination-mediated proteasomal degradation. Upon Wnt stimulation, β-catenin translocates into the nucleus, interacts with the nuclear transcription factor TCF/LEF and transactivates downstream genes[26, 27]. Based on these findings, we explored the effect of WSB1 on the cellular location and ubiquitination of β-catenin.

**Figure 3.**
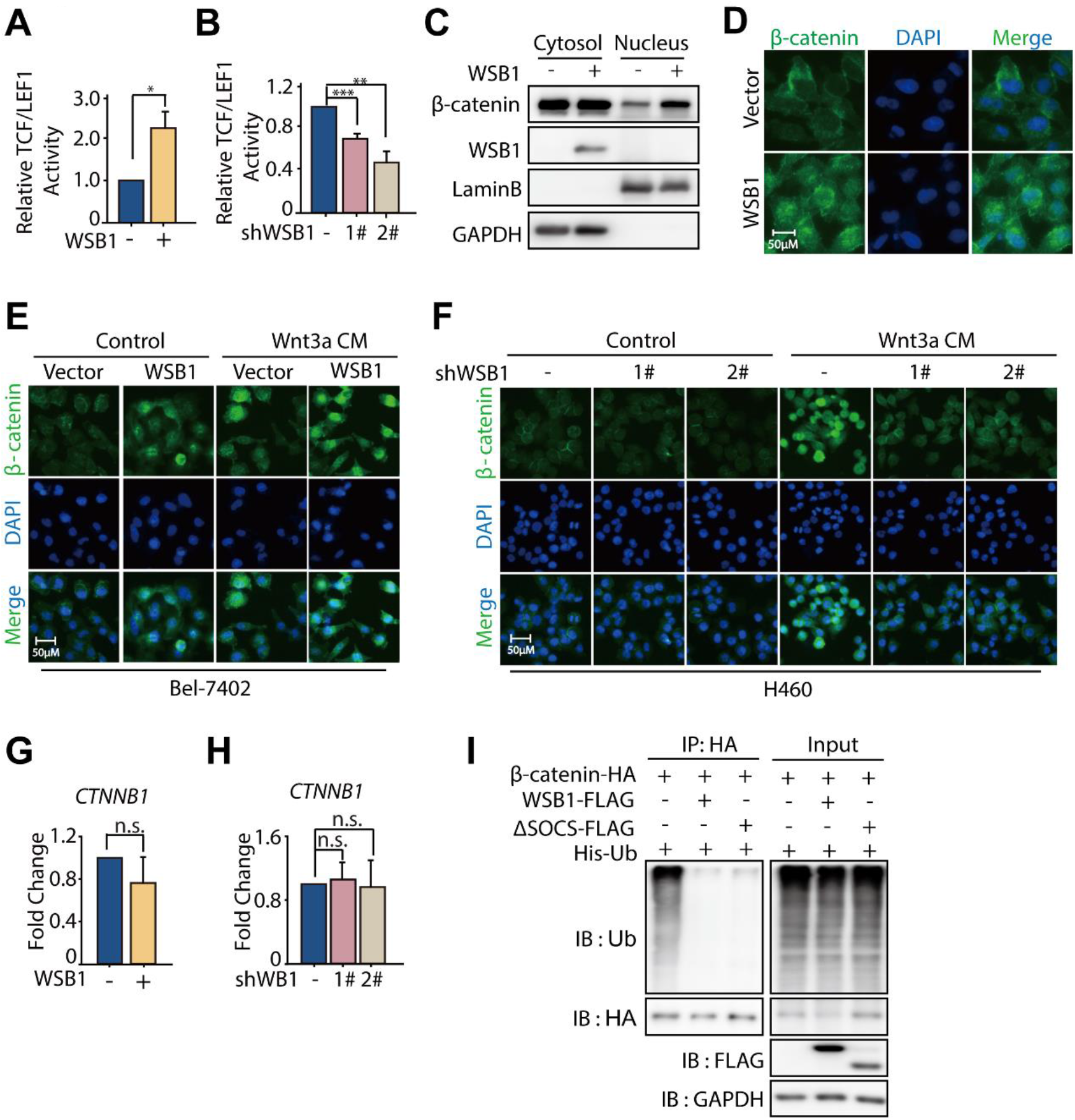
WSB1 promotes β-catenin nucleus translocation and inhibits its ubiquitination. (A) Bel-7402 cells were co-transfected with vector or pCDH-WSB1 and pGL4.14-TCF/LEF1-luciferase for 48 hours. Luciferase activity was normalized by cell numbers and shown as fold change. Data was represented as the means ± SD, n = 3; Statistical significance was determined by Student’s t-test. *, p < 0.05 (versus vector group). (B) H460 cells were transduced with lentivirus WSB1 shRNA followed by transfection with vector or pCDH-WSB1 and TCF/LEF1-luciferase for 48 hours. Luciferase activity was normalized by cell numbers and shown as fold change. Data was represented as the means ± SD, n = 3; Statistical significance was determined by Student’s t-test. **, p < 0.01; ***, p < 0.001 (versus control group). (C) Bel-7402 cells were infected with lentivirus pCDH Vector or lentivirus pCDH-WSB1 for 72 hours and collected for cell fraction (nuclei and cytoplasm) separation. Cellular localization of β-catenin was detected by Western blotting in cell fraction. GAPDH was used as loading control for cytoplasm fraction and Lamin B was for nucleus fraction. (D) Immunofluorescence staining of β-catenin in Bel-7402 cells infected with lentivirus pCDH Vector or lentivirus pCDH-WSB1. (E) Bel-7402 cells infected with lentivirus pCDH Vector or lentivirus pCDH-WSB1 were treated with Wnt3a condition medium (Wnt3a CM) for 16 hours, and stained with β-catenin antibody by immunofluorescence. (F) H460 cells infected with lentivirus Vector or lentivirus WSB1 shRNAs were treated with Wnt3a condition medium (Wnt3a CM) for 16 hours, and stained with β-catenin antibody by immunofluorescence. (G) Relative mRNA level of *WSB1* and *CTNNB1* in Bel-7402 cells infected with lentivirus pCDH-WSB1 or 72 hours. (H) Relative mRNA level of *WSB1* and *CTNNB1* in H460 cells infected with lentivirus WSB1 shRNA for 72 hours. Data was represented as the means ± SD, n = 3; Statistical significance was determined by Student’s t-test. n.s., no significant difference (p>0.05). (I) 293FT cells were infected with lentivirus pCDH vector or lentivirus pCDH -WSB1 or lentivirus pCDH-ΔSOCS for 72 hours. Then the cells were plated and transfected with pCDNA-3.0-β-catenin-HA and His-Ub for 48 hours and treated with MG132 (10 μM) for 8 hours before collecting cells. Then the cells were lysed and for immunoprecipitated with anti-HA antibody followed by immunoblotting with anti-ubiquitination and anti-HA antibody.

First, we determined the effect of WSB1 on the nuclear translocation of β-catenin by performing a nucleocytoplasmic separation assay. As shown in Figure 3C, WSB1 overexpression increased nuclear β-catenin expression, while cytoplasmic β-catenin was not significantly changed, suggesting a promoting effect of WSB1 on β-catenin nuclear translocation. Similarly, an immunofluorescence assay in Bel-7402 cells also proved that WSB1 overexpression could promote β-catenin entry into the nucleus compared that in to the vector control group (Figure 3D). Given that Wnt signalling could stabilize β-catenin and promote its nuclear translocation, we next tested the effect of WSB1 on β-catenin nuclear translocation under Wnt stimulation conditions. As expected, Bel-7402 cells treated with Wnt3a-conditioned medium (Wnt3a-CM) markedly induced β-catenin nuclear translocation (Figure 3E). Interestingly, WSB1 overexpression further promoted the nuclear translocation of β-catenin in coordination with Wnt3a (Figure 3E). Moreover, knocking down WSB1 in H460 cells abolished the promoting effect of Wnt3a-conditioned medium (Figure 3F). Therefore, these data clearly suggested that WSB1 promoted β-catenin nuclear translocation under both basal and Wnt stimulation conditions.

To further address the role of WSB1 on the β-catenin ubiquitination status, RT-PCR was performed to investigate whether WSB1 influenced the mRNA level of *CTNNB1* (the gene encoding β-catenin). As shown in Figure 3G and 3H, manipulating the expression of WSB1 in Bel-7402 and H460 cells had no influence on *CTNNB1*. Together with the fact that WSB1 increased the expression of β-catenin (Figure 2D and 2E), these results further indicated that WSB1 regulated β-catenin at the posttranscriptional level. Subsequently, we investigated whether WSB1 participated in the ubiquitination process of β-catenin. As illustrated in Figure 3I, wild-type WSB1, as well as ΔSOCS WSB1, inhibited the ubiquitination of β-catenin. Thus, our data further suggested that WSB1 significantly inhibited the ubiquitination of β-catenin, which resulted in β-catenin protein stabilization and nuclear translocation.

### 4. WSB1 interacts with the destruction complex through the scaffold protein AXIN1

It is known that in the absence of Wnt, β-catenin is captured by the destruction complex composed of AXIN1, APC, casein kinase 1 (CK1) and glycogen synthase kinase 3β (GSK3β), subsequently losing the ability to enter the nucleus and being recognized by its E3 ligase adaptor β-TRCP[14] (Figure 4A). Given that (i) WSB1 decreased the ubiquitination of β-catenin (Figure 3I) and (ii) co-immunoprecipitation experiments indicated that both wild-type and ΔSOCS WSB1 could interact with the four major exogenous and endogenous components (AXIN1, CK1, GSK3β, and β-TRCP) of the β-catenin destruction complex (Figure 4B and 4C), we wanted to investigate how WSB1 modulated the β-catenin destruction complex. Thus, we first checked whether WSB1 has an effect on the protein expression of the major components of the destruction complex. As shown in Figure 4D, overexpressing wild-type and ΔSOCS WSB1 did not inhibit AXIN1, CK1, GSK3β, or β-TRCP expression in either 293FT or Bel-7402 cells, which roughly ruled out the possibility of WSB1-driven inhibition of destruction complex components.

**Figure 4.**
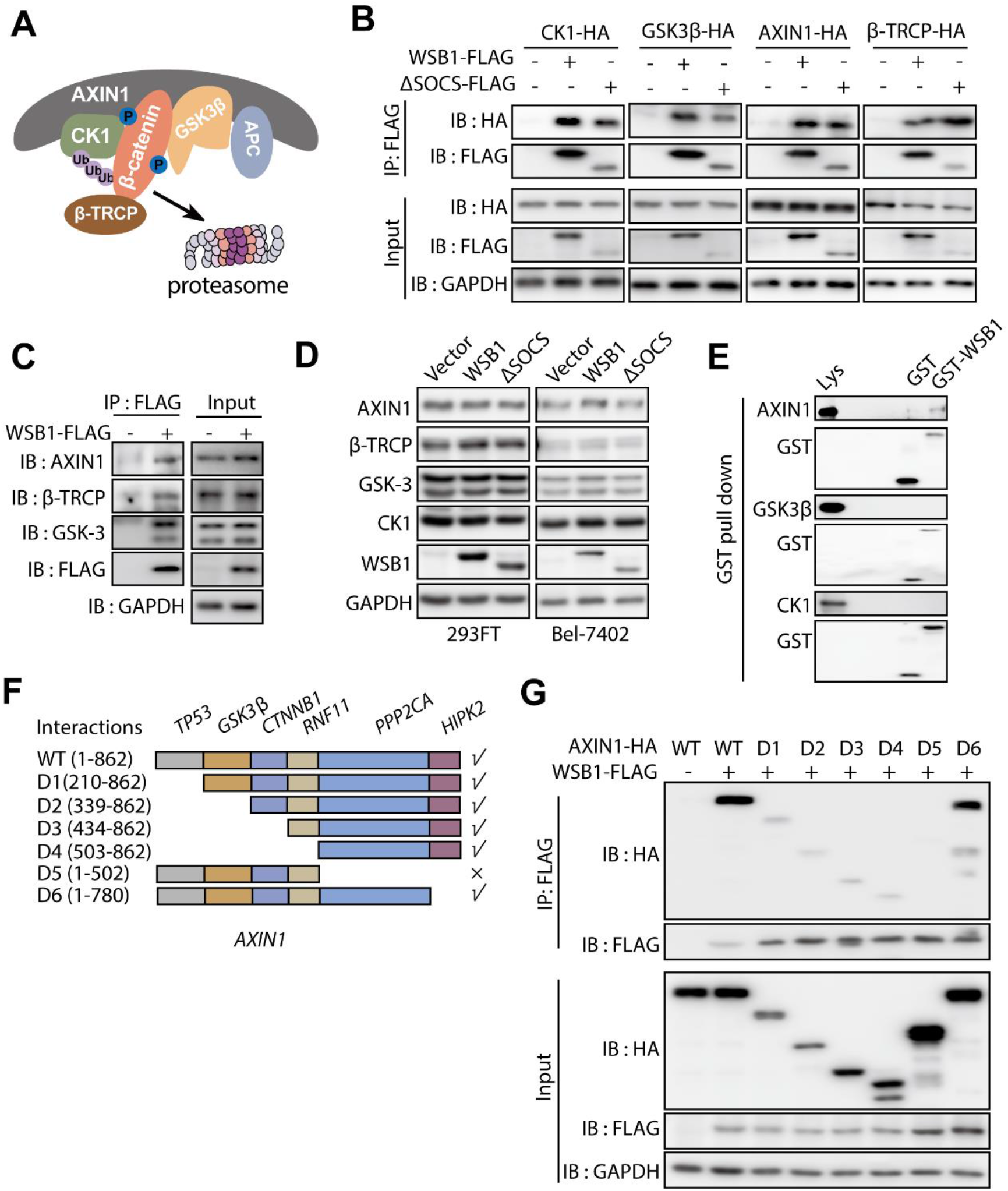
WSB1 interacts with the destruction complex through the scaffold protein AXIN1. (A) Schematic representation of the β-catenin destruction complex. (B) 293FT cells were transfected with β-catenin destruction complex components plasmids (pCDNA-3.0-CK1-HA or pCDNA-3.0-GSK3β-HA or pCDNA-3.0-AXIN1-HA or pCDNA-3.0-β-TRCP-HA) or co-transfected with pCDH-WSB1-FLAG or pCDH-ΔSOCS-FLAG for 48 hours. Then the cells were lysed and for immunoprecipitated with anti-FLAG antibody followed by immunoblotting with anti-HA antibody. (C) 293FT cells were transfected with pCDH vector or pCDH-WSB1-FLAG for 48 hours. Then the cells were lysed and for immunoprecipitated with anti-FLAG antibody followed by immunoblotting with anti-AXIN1 or anti-β-TRCP or anti-GSK3 or anti-FLAG antibody. (D) Western blotting of AXIN1, β-TRCP, GSK3β and CK1 in 293FT cells and Bel-7402 cells infected with lentivirus pCDH vector or lentivirus pCDH-WSB1 or lentivirus pCDH-ΔSOCS for 72 hours. (E) 293FT cells were transfected with pCDNA-3.0-AXIN1-HA or pCDNA-3.0-CK1-HA or pCDNA-3.0-GSK3β-HA for 48 hours. The cells were lysed by 4% SDS lysis buffer and then diluted in PBS buffer to make the SDS concentration was 0.1%. 5 μg GST protein or WSB1-GST fusion protein were incubated with 10 ul GST-seflinose™ resin for 2 hours at 4°C followed by incubating with the lysates overnight at 4°C and immunoblotting with anti-HA antibody and anti-GST antibody. (F) Schematic representation of AXIN1 domains and a series of AXIN1-HA deletion mutations. (G) 293FT cells were transfected with pCDH-WSB1-FLAG and a series of AXIN1-HA deletion mutations for 48 hours. Then the cells were lysed and for immunoprecipitated with anti-FLAG antibody followed by immunoblotting with anti-HA antibody.

Interestingly, when we incubated the recombinant GST-tagged WSB1 with cell lysate to identify the potential direct interaction proteins, the results indicated that AXIN1 directly interacted with WSB1, while the other complex components did not (Figure 4E). Given that AXIN1 is the scaffold protein in the destruction complex, acting as the recruiter of the other proteins[28, 29], we further hypothesized that AXIN1 played a critical role in the interaction between destruction complex components and WSB1. To further determine the regions mediating the interaction between WSB1 and AXIN1, we constructed a series of truncated AXIN1 isoforms and tested their interactions with WSB1 (Figure 4F). As shown in Figure 4G, we found that the D5 region truncation of AXIN1 (Δ503-862) abolished the interaction of WSB1 and AXIN1. Collectively, these data implied that WSB1 directly interacted with the destruction complex through the scaffold protein AXIN1, and the 503-780 region of AXIN1 was responsible for this interaction.

### 5. WSB1 affects destruction complex-PPP2CA assembly and E3 ubiquitin ligase adaptor β-TRCP recruitment

Previous reports have suggested that the 503-582 region of AXIN1 is the binding site of PPP2CA, which is the catalytic subunit of the serine/threonine protein phosphatase 2A (PP2A)[30]. As PP2A can dephosphorylate the β-catenin destruction complex and affect the stability of β-catenin as well as its downstream genes[31, 32], we explored whether the interaction of AXIN1 and WSB1 regulated the recruitment of PPP2CA and its function. Co-immunoprecipitation assays were carried out in 293FT cells overexpressing WSB1, AXIN1 and PPP2CA, and the results indicated that WSB1 overexpression promoted the interaction between AXIN1 and PPP2CA (Figure 5A and 5B). Importantly, the promoting effect of WSB1 on AXIN1-PPP2CA complex formation was dose-dependent (Figure 5C and 5D).

**Figure 5.**
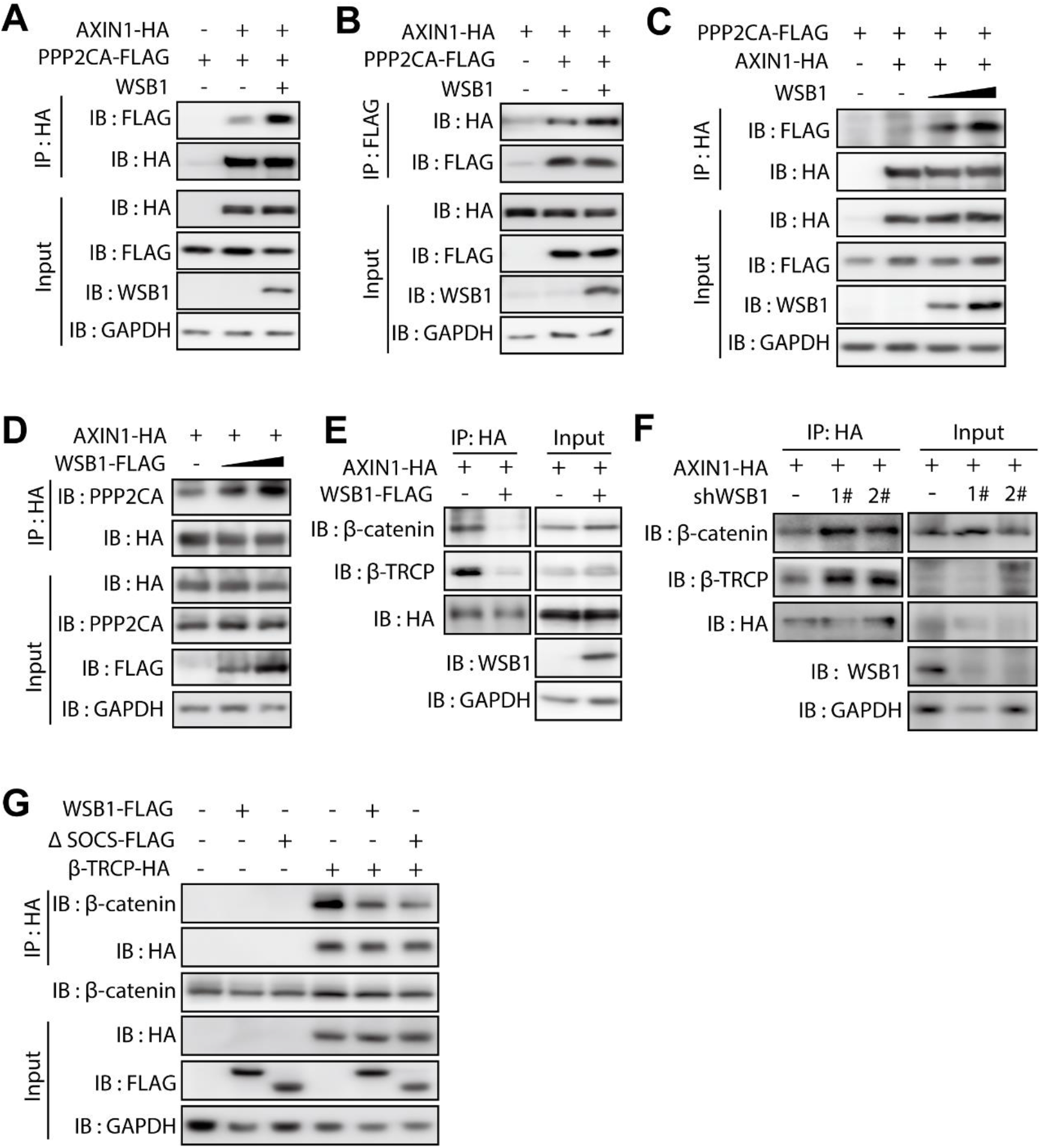
WSB1 affects the destruction complex-PPP2CA assembly and the E3 ubiquitin ligase adaptor β-TRCP recruitment. (A) 293FT cells were transfected with pCDNA3.0-PPP2CA-FLAG, or co-transfected with pCDNA3.0-PPP2CA-FLAG and pCDNA3.0-AXIN1-HA or co-transfected with pCDNA3.0-PPP2CA-FLAG, pCDNA3.0-AXIN1-HA and pCDH-WSB1-FLAG for 48 hours. Then the cells were lysed and for immunoprecipitated with anti-FLAG antibody followed by immunoblotting with anti-HA antibody or immunoprecipitated with anti-HA antibody followed by immunoblotting with anti-FLAG antibody. (B) 293FT cells were transfected with pCDNA3.0-AXIN1-HA, or co-transfected with pCDNA3.0-AXIN1-HA and pCDNA3.0-PPP2CA-FLAG or co-transfected with pCDNA3.0-PPP2CA-FLAG, pCDNA3.0-AXIN1-HA and pCDH-WSB1 for 48 hours. Then the cells were lysed and for immunoprecipitated with anti-FLAG antibody followed by immunoblotting with anti-HA antibody or immunoprecipitated with anti-HA antibody followed by immunoblotting with anti-FLAG antibody. (C) 293FT cells were transfected with pCDNA3.0-PPP2CA-FLAG, or co-transfected with pCDNA3.0-PPP2CA-FLAG and pCDNA3.0-AXIN1-HA, or co-transfected with pCDNA3.0-PPP2CA-FLAG, pCDNA3.0-AXIN1-HA and different doses of pCDH-WSB1(1 μg or 2 μg) for 48 hours. Then the cells were lysed and for immunoprecipitated with anti-HA antibody followed by immunoblotting with anti-FLAG antibody. (D) 293FT cells were transfected with pCDNA3.0-AXIN1-HA, or co-transfected with pCDNA3.0-AXIN1-HA and different doses of pCDH-WSB1-FLAG (1 μg or 2 μg) for 48 hours. Then the cells were lysed and for immunoprecipitated with anti-HA antibody followed by immunoblotting with anti-PPP2CA antibody. (E) 293FT cells were infected with lentivirus pCDH vector or lentivirus pCDH -WSB1 for 72 hours. Then the cells were lysed and for immunoprecipitated with anti-HA antibody followed by immunoblotting with anti-β-catenin, anti-β-TRCP and anti HA antibody. (F) 293FT cells were infected with lentivirus shRNA vector or lentivirus WSB1 shRNA for 72 hours. Then the cells were plated and transfected with pCDNA-3.0-AXIN1-HA for 48 hours. Then the cells were lysed and for immunoprecipitated with anti-HA antibody followed by immunoblotting with anti-β-catenin, anti-β-TRCP and anti HA antibody.(G) 293FT cells were infected with lentivirus pCDH vector or lentivirus pCDH -WSB1 or lentivirus pCDH-ΔSOCS for 72 hours. Then the cells were plated and transfected with pCDHA3.0 vector or pCDNA-3.0-β-TRCP-HA for 48 hours. Then the cells were lysed and for immunoprecipitated with anti-HA antibody followed by immunoblotting with anti-β-catenin and anti-HA antibody.

Although the interaction between PPP2CA and AXIN1 is required for the proper function of the β-catenin destruction complex, recent evidence further suggests that the dynamic interaction of AXIN1 and PPP2CA, rather than a steady-state AXIN1-PPP2CA complex formation, is pivotal in maintaining the β-catenin destruction complex function[26]. Thus, we wondered whether the interaction between AXIN1 and PPP2CA promoted by WSB1 would regulate the assembly and function of the β-catenin destruction complex. The results showed that overexpressing WSB1 in 293FT cells significantly inhibited the interaction between β-catenin and AXIN1 as well as β-TRCP and AXIN1 (Figure 5E), while knocking down WSB1 increased the interactions (Figure 5F), which was consistent with the results of WSB1 overexpression. To further clarify the function of WSB1 in β-catenin stability regulation, we investigated whether WSB1 affected the interaction between β-catenin and its E3 ligase adaptor β-TRCP. As shown in Figure 5G, both wild-type and ΔSOCS WSB1 inhibited the interaction between β-catenin and β-TRCP. Taken together, our data showed that WSB1 inhibited destruction complex assembly and E3 ubiquitin ligase β-TRCP recruitment and therefore promoted the stability of β-catenin.

### 6. The WSB1/c-Myc feedforward circuit participates in the development of cancer

Next, we explored the physiological relevance of the WSB1/c-Myc feedforward circuit. Because the c-Myc and the Wnt/β-catenin pathway both play important roles in tumour initiation and tumour stemness regulation, we conducted colony formation and sphere formation assays to study whether the WSB1/c-Myc feedforward circuit influenced cell proliferation and stemness. The results indicated that overexpressing WSB1 by lentivirus infection of Bel-7402 cells increased the colony numbers, and when co-overexpressed with c-Myc, WSB1 further promoted colony formation compared to WSB1 overexpression alone (Figure S3A and S3B). To further validate this result, we knocked down WSB1 in H460 cells under both basal conditions and c-Myc overexpression conditions. As expected, the overexpression of c-Myc significantly increased colony formation in H460 cells (Figure S3C). Of note, WSB1 inhibition significantly inhibited the increase in colony formation induced by c-Myc (Figure S3C and S3D). These results suggest that the WSB1/c-Myc feedforward circuit promoted cancer cell proliferation.

To examine the effect of WSB1 on cell stemness, a sphere formation assay was performed in Bel-7402 cells. As shown in Figure 6A and 6B, both wild-type and ΔSOCS WSB1 overexpression significantly increased the cell sphere numbers. When c-Myc was also overexpressed, WSB1 further promoted sphere formation (Figure 6C and 6D). A similar effect was also observed in another HCC cell line, HuH7 (Figure S3E and S3F), suggesting that WSB1 played a critical role in cell stemness.

**Figure 6.**
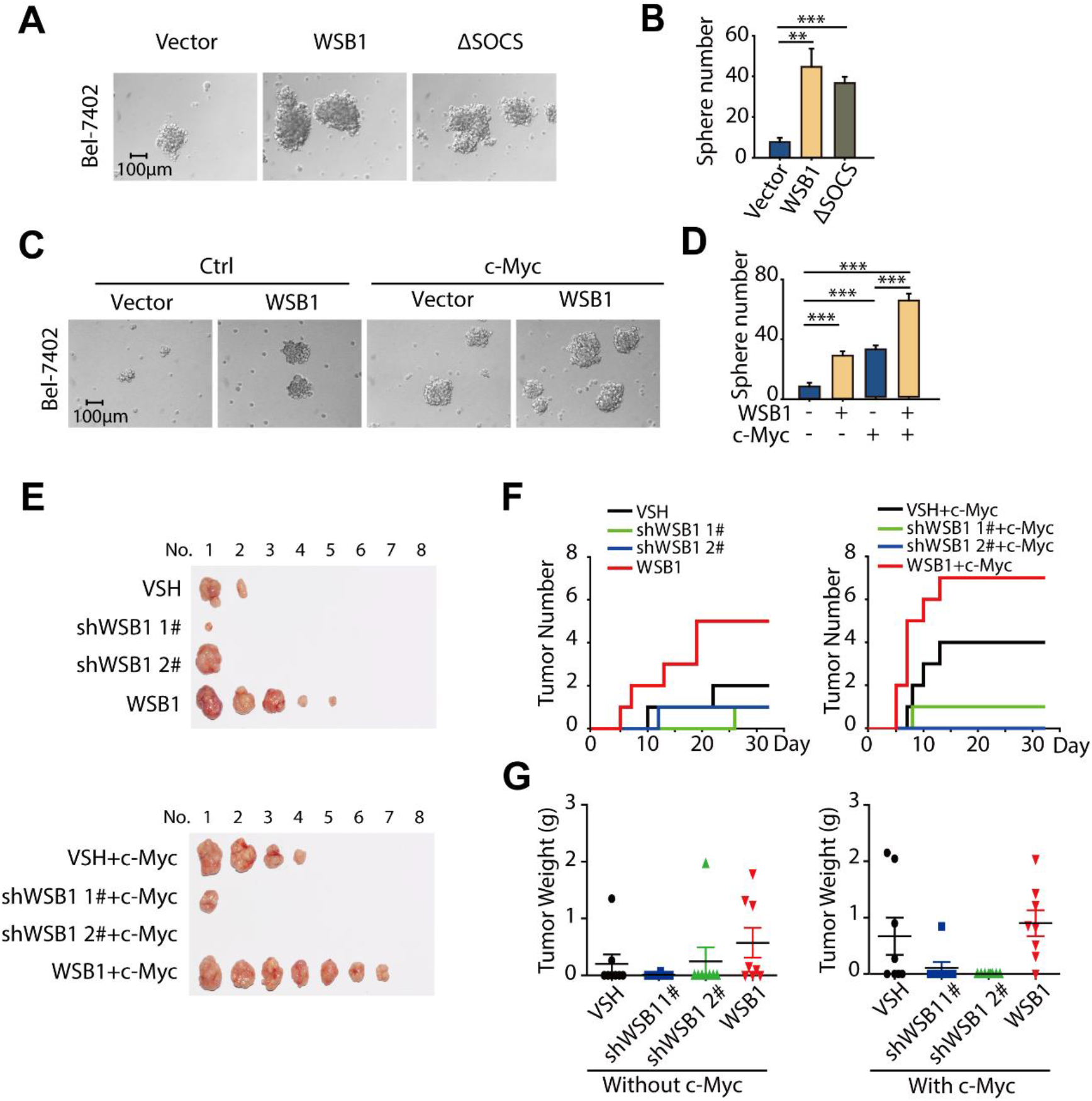
WSB1/c-Myc feedforward circuit participated in the development of cancer. (A) Sphere formation assay was performed in Bel-7402 cells transduced with lentivirus pCDH or pCDH-WSB1 or pCDH-ΔSOCS. Representative photographs were taken by microscope. (B) Statistical analysis of cell colonies. Data was represented as the means ± SD, n = 3; Statistical significance was determined by Student’s t-test. **, p < 0.01; ***, p < 0.001 (versus vector group). (C) Sphere formation assay was performed in Bel-7402 cells transduced with lentivirus pCDH or pCDH-WSB1 or pCDH-c-Myc or both. Representative photographs were taken by microscope. (D) Statistical analysis of cell colonies. Data was represented as the means ± SD, n = 3; Statistical significance was determined by Student’s t-test. ***, p < 0.001 (versus vector group). (E) Schematic illustration of experimental design. Mice transplanted with different group of H460 cells were observed every day, and tumor numbers and growing periods of mice were recorded. (F) Photographs of tumors from different groups. In the end of the experiment, the mice were sacrificed and tumors were dissected. (G and H) Analysis of tumor numbers and tumor weights in different groups. The average tumor weights of each group were presented as mean ± SEM, n=8.

Encouraged by the in cell results, we wondered whether this effect could be recapitulated *in vivo*. Thus, we performed an H460 cell xenograft tumour assay in nude mice, and 8 groups of cells (VSH and VSH+c-Myc; shWSB1 #1 and shWSB1 #1+c-Myc; shWSB1 #2 and shWSB1 #2+c-Myc; WSB1 and WSB1+c-Myc) were subcutaneously injected into the same mice. Cells without c-Myc overexpression were injected into the left side, while cells with c-Myc overexpression were injected into the right side. The tumour formation time and tumour growth were monitored. When the end of the experiment was reached, the mice were sacrificed, and tumours were dissected. As shown in Figure 6E in line with the cell results, overexpressing WSB1 significantly promoted tumour formation (2 tumours in the VSH group *vs*. 5 tumours in the WSB1 group). Moreover, WSB1 overexpression further increased tumour formation compared to c-Myc single overexpression (4 tumours in the VSH+c-Myc group *vs*. 7 tumours in the WSB1+c-Myc group), while knocking down WSB1 completely abolished the c-Myc promoting effect (1 tumour in the shWSB1 #1+c-Myc group and 0 tumours in the shWSB1 #2+c-Myc group) (Figure 6E). Tumour formation curves and tumour weight curves also showed that WSB1 could improve the tumour promoting effect induced by c-Myc (Figure 6F and 6G).

To validate the finding that the WSB1/c-Myc feedforward circuit is present in tumours, immunohistochemistry experiments were subsequently performed to examine the expression of WSB1, c-Myc and β-catenin. As shown in Figure S3G, WSB1 overexpression was accompanied by relatively high levels of c-Myc and β-catenin. When WSB1 was silenced in the c-Myc-overexpressing group, the protein level of c-Myc was remarkably decreased compared with that of the c-Myc-overexpressing group (Figure S3G). Moreover, overexpressing WSB1 further increased the c-Myc protein level (Figure S3G). These data further proved that the WSB1/c-Myc feedforward circuit existed *in vivo* and participated in the development of cancer.

## Discussion

c-Myc is documented to be related not only to the initiation of many malignant tumours but also to malignant progression, such as tumour resistance and recurrence[7]. Apart from its nuclear location, the amplification or transcriptional dysregulation of c-Myc is accompanied by an anabolic transcriptional response that promotes the metabolic adaptation of cancer cells, both of which limit the development of cancer therapy strategies directly targeting c-Myc[33, 34]. Studies have implied that the regulation of c-Myc is mainly characterized by transcription and protein stability. The half-life of c-Myc is approximately 20-30 minutes, indicating that the protein level of c-Myc is changed dynamically in cellular activity, which means that it is difficult to target[35, 36]. In contrast, as the transcription of c-Myc is regulated by multiple signalling pathways, including the classic Wnt/β-catenin pathway[14, 37], the upstream regulation of c-Myc through β-catenin would provide a critical theoretical basis for c-Myc targeting strategies. In this regard, our results suggest that the WSB1/c-Myc feedforward circuit plays an important role in tumorigenesis, and WSB1 inhibition can effectively block the tumour-promoting effect of c-Myc.

Composed of seven WD40 repeat domains and a SOCS box responsible for Elongin B/C binding and substate recognition, WSB1 has been reported to regulate the degradation of multiple substrates, all of which are based on its E3 enzyme activity. However, in our study, we found that WSB1 could enhance the expression of c-Myc even when it lacked the SOCS box, which means that the promoting effect of WSB1 is independent of its E3 ligase activity. As an adaptor of the E3 ligase, WSB1 has other biological functions in addition to mediating substrate recognition of the ubiquitination process. From this perspective, our findings propose for the first time a new role for WSB1 in cancer development, which enriches the theoretical knowledge of WSB1 and the ubiquitin proteasome system. Although the WSB1 inhibitor could be ideally designed based on the interaction between WSB1 and its substrates, our results further suggest that there are other functional domains regulating tumour progression. Thus, it might be worth further developing a PROTAC (proteolysis targeting chimaera) molecule to degrade the WSB1 protein, rather than a protein-protein binding inhibitor, to fully inhibit its tumour promoting effect.

Many studies have reported that c-Myc can form feedback loops in tumour regulation; for example, c-Myc can directly transactivate EBNA1-binding protein 2 (EBP2), while EBP2 affects F-box/WD repeat-containing protein 7 (FBW7), the E3 ligase of c-Myc, to inhibit c-Myc degradation, which ultimately promotes cancer progression[38, 39]. Moreover, c-Myc can form a feedback loop with its downstream gene Eukaryotic initiation factor 4F (eIF4F) to promote the cell proliferation induced by c-Myc[24]. In our study, we found that WSB1 forms a feedforward circuit with c-Myc through the Wnt/β-catenin pathway to significantly promote cancer development, and at the same time, c-Myc can transactivate WSB1 to form a circuit loop. Thus, the WSB1/c-Myc feedforward circuit, which can also be considered as a β-catenin/c-Myc loop, provides new insight into the relationship between β-catenin and c-Myc. Moreover, β-catenin and c-Myc have been reported to be associated with cancer stemness in several studies[40, 41], which endows the β-catenin/c-Myc loop with stemness-promoting ability compared with other reported feedback loops.

However, there are some limitations to our mechanistic study, as the precise regulatory role and the direct substrate protein of PPP2CA in this feedback loop are still unclear and remain to be further explored in future studies. In addition, numerous studies have pointed out that the phosphorylation and dephosphorylation of the destruction complex, especially of AXIN1, precisely control β-catenin stability and Wnt pathway activation[42]. The phosphatases PP2A and PP1 are both reported to be correlated with the complex; of note, PP2A can interact with both AXIN1 and APC, while PP1 only binds with AXIN1[30, 42, 43]. These facts indicate that it is necessary to explore whether the promotional effects of WSB1 on PP2A are specific to the WSB1/c-Myc loop; in other words, determine if other phosphatases also participate in the regulation of WSB1. Moreover, as PP1 is reported to dephosphorylate AXIN1 and induce a conformational switch[42], it is interesting to determine whether different structural characteristics and assembly statuses of AXIN1 play a critical role in WSB1 regulation. More importantly, a recent study proved that Myc regulated the immune response through PD-L1[6], which greatly aroused our interest to explore whether WSB1 could affect PD-L1 and immune therapy through the WSB1/c-Myc feedforward loop.

## Conclusions

In summary, our results prove that WSB1 is a direct target gene of c-Myc and can form a feedforward circuit with c-Myc through β-catenin, which leads to cancer initiation and development. This finding not only provides new targeting strategies for c-Myc in cancer therapy but also enriches the c-Myc and WSB1 crosstalk network theory. The discovery of the WSB1/β-catenin/c-Myc axis clarifies the critical role of WSB1 in the Wnt/β-catenin signalling pathway, which also provides new possibilities for indirectly intervening in the β-catenin pathway in cancer treatment.

## List of abbreviations

TFs: Transcription factors
WSB1: WD repeat and SOCS box containing 1
HIF1-α: Hypoxia induced factor 1-alpha
RhoGDI2: Rho GDP dissociation inhibitor 2
HCC: Hepatocellular carcinoma
Wnt3a-CM: Wnt3a-conditioned medium
CK1: Casein kinase 1
GSK3β: Glycogen synthase kinase 3β
EBP2: EBNA1-binding protein 2
FBW7: F-box/WD repeat-containing protein 7
eIF4F: Eukaryotic initiation factor 4F

## Declarations

### Ethics approval and consent to participate

Maintenance and experimental procedures for the mice studies were approved by Zhejiang University’s IACUC (IACUC-s19-038).

### Consent for publication

Not applicable.

### Availability of data and materials

The datasets generated and/or analysed during the current study are not publicly available but are available from the corresponding author on reasonable request.

### Competing interests

No conflict of interest exists in the submission of this manuscript.

### Funding

This work was supported by grants from Zhejiang Provincial Natural Science Foundation (No. Y18H310001 to Ji Cao), the National Natural Science Foundation of China (No. 81872885 to Ji Cao; No.81625024 to Bo Yang), and the Talent Project of Zhejiang Association for Science and Technology (No.2018YCGC002 to Ji Cao).

### Authors’ contributions

Ji Cao, Yanling Gong, Xiaomeng Gao, and Bo Yang designed the research; Ji Cao, Xiaomeng Gao, and Bo Yang wrote the manuscript; Yanling Gong, Xiaomeng Gao, Jieqiong You, Meng Yuan, Liang Fang, and Haiying Zhu performed the biochemical and cellular studies; Yanling Gong, Xiaomeng Gao, and Jieqiong You conducted the animal studies; Ji Cao, Yanling Gong, Xiaomeng Gao analyzed the results; Ji Cao, Hong Zhu, Meidan Ying, Qiaojun He and Bo Yang directed the study.

## Acknowledgements

Not applicable.

---

**Figure S1.**
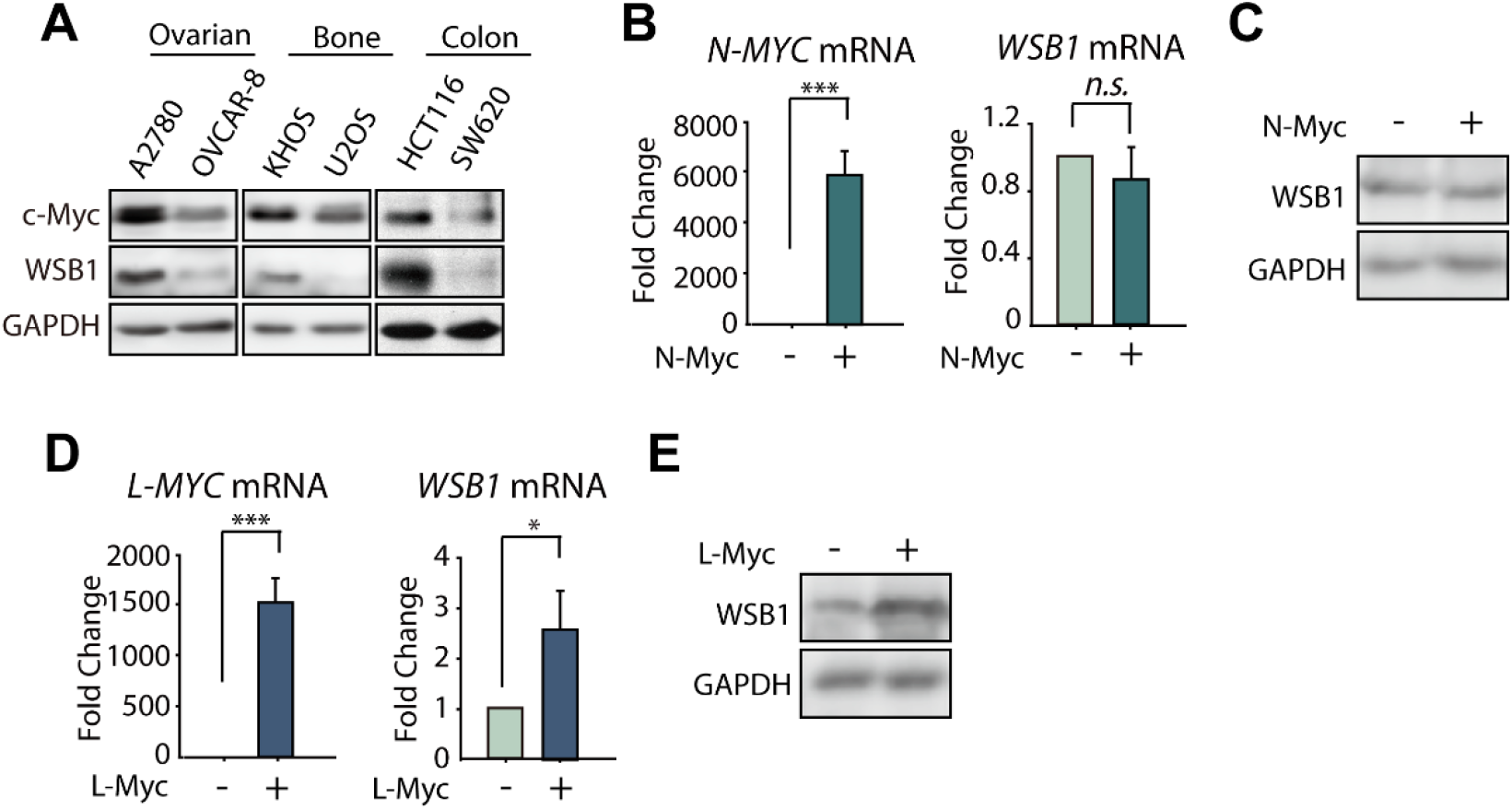
(A) Western blotting of WSB1 and c-Myc expression in different tumor cell lines from various tissues. (B) Relative mRNA levels of *WSB1* and *N-MYC* were detected after transfecting with pCDH-N-Myc plasmids for 48 hours. Data was represented as the means ± SD, n = 3; Statistical significance was determined by Student’s t-test. n.s., no significant difference (p>0.05); ***, p < 0.001;(versus vector group). (C) Wstern blotting of WSB1 after transfecting with pCDH-N-Myc plasmids for 48 hours. (D) Relative mRNA levels of *WSB1* and *L-MYC* were detected after transfecting with pCDH-L-Myc plasmids for 48 hours. Data was represented as the means ± SD, n = 3; Statistical significance was determined by Student’s t-test. *, p < 0.05; ***, p < 0.001;(versus vector group). (E) Wstern blotting of WSB1 after transfecting with pCDH-L-Myc plasmids for 48 hours.

**Figure S2.**
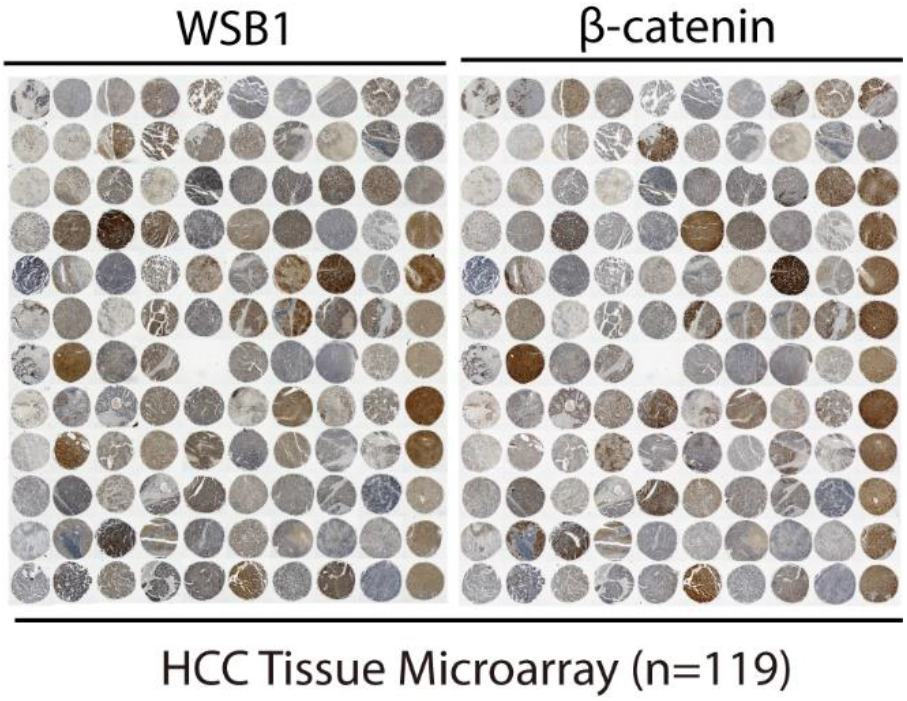
Immunohistochemical (IHC) staining of β-catenin in hepatocellular carcinoma tissue. Here, the WSB1 results are from Figure.1B.

**Figure S3.**
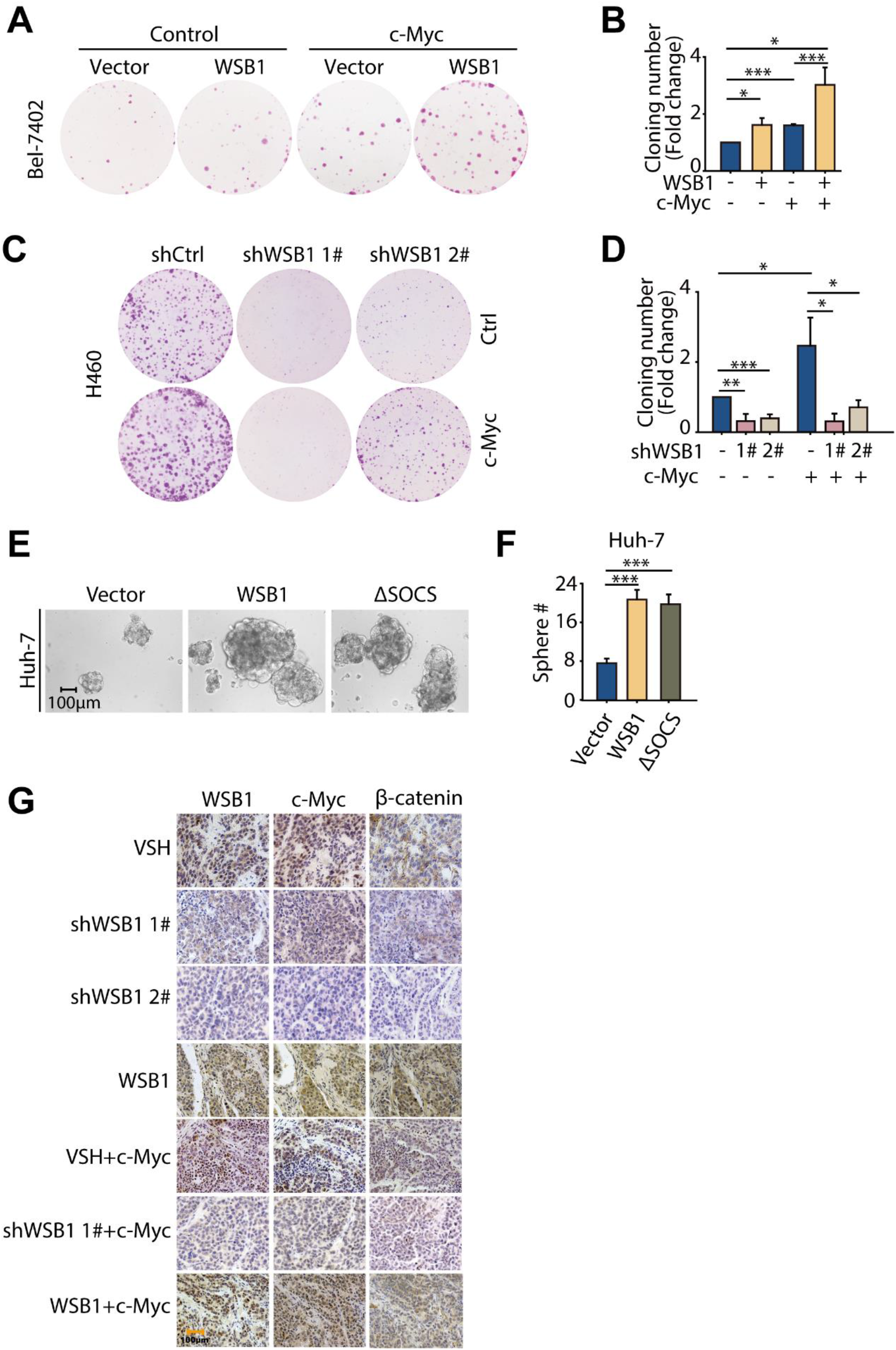
(A) Cloning formation assay was performed in Bel-7402 cells transduced with lentivirus pCDH or pCDH-WSB1 or pCDH-c-Myc or both. Cells were stained with the sulforhodamine B and photographed by E-Gel Imager. (B) Cloning numbers were counted and shown as fold change. Data was represented as the means ± SD, n = 3; Statistical significance was determined by Student’s t-test. *, p < 0.05; ***, p < 0.001 (versus vector group). (C) Cloning formation assay was performed in H460 cells transduced with lentivirus shControl or shWSB1 or pCDH-c-Myc or both. Cells were stained with the sulforhodamine B and photographed by E-Gel Imager. (D) Cloning numbers were counted and shown as fold change. Data was represented as the means ± SD, n = 3; Statistical significance was determined by Student’s t-test. *, p < 0.05; **, p < 0.01; ***; p < 0.001 (versus control group). (E) Sphere formation assay was performed in HuH-7 cells transduced with lentivirus pCDH or pCDH-WSB1 or pCDH-ΔSOCS. Representative Photographs were taken by microscope. (F) Statistical analysis of cell colonies. Data was represented as the means ± SD, n = 3; Statistical significance was determined by Student’s t-test. ***, p < 0.001 (versus vector group). (G) Sacrificed the nude mice at the end of the xenograft experiment and dissected the tumor. Immunohistochemical (IHC) staining of WSB1, c-Myc and β-catenin in tumor tissue from different groups.

**Table S1. RNA-seq results**

## Notes

### Competing Interest Statement

The authors have declared no competing interest.

### Summary of Updates

1. For the reason of the network interruption previously, figures from 4-S3 were missing in the uploading process. These figures are uploaded again to fullfill this article. 2. Besides, the order of the corresponding author has a miner change. 3. Supplemental files updated.

